# Mapping the impacts of hurricanes Maria and Irma on banana production area in the Dominican Republic

**DOI:** 10.1101/2020.09.20.304899

**Authors:** Varun Varma, William Thompson, Solhanlle Bonilla Duarte, Pius Krütli, Johan Six, Daniel P Bebber

## Abstract

Extreme weather events can have devastating impacts on agricultural systems, and the livelihoods that depend on them. Tools for rapid, comprehensive and cost-effective assessment of impacts, especially if carried out remotely, can be of great value in planning systematic recovery of production, as well as assessing risks from future events. Here, we use openly available remote sensing data to quantify the impacts of hurricanes Irma and Maria in 2017 on banana production area in the Dominican Republic — the world’s largest producer of organic bananas. Further, we assess the risk to current production area if a similar extreme event were to re-occur. Hurricane associated damage was mapped using a simple change detection algorithm applied to Synthetic Aperture Radar (SAR) data over the three main banana growing provinces of northern Dominican Republic, i.e. Monte Cristi, Valverde and Santiago. The map of hurricane affected area was overlaid with banana plantation distributions for 2017 and 2019 that were mapped (accuracy = 99.8%) using a random forest classifier, and a combination of SAR and multi-spectral satellite data. Our results show that 11.35% of banana plantation area was affected by hurricane damage in 2017. Between 2017 and 2019, there was a high turnover of plantation area, but with a net gain of 10.8%. However, over a quarter (26.9%) of new plantation area spatially overlapped with regions which had seen flooding or damage from hurricanes in 2017. Our results indicate that banana production systems in northern Dominican Republic saw extensive damage in the aftermath of hurricanes Irma and Maria. While production area has recovered since then, a substantial proportion of new plantations, and a greater fraction of production area in general, occur at locations at risk from future extreme events.

## Introduction

Extreme weather events, such as floods and droughts, can have a considerable impact on agricultural production systems across the globe by reducing agricultural output (IPCC, 2012; Lesk, Rowhani & Ramankutty, 2013). The impacts of these events are felt in terms of reducing local and regional food security, and can also have significant economic consequences via their negative influence on income generation and livelihoods (Wheeler & Braun, 2013). With climate change likely to increase the frequency and severity of extreme weather (IPCC, 2014), quantifying the extent of damage after such events is valuable information towards developing required risk management and mitigation strategies. However, in the immediate aftermath of pulsed and high intensity events, such as flooding following hurricanes, ensuring human safety, health and rehabilitation take precedence over physical agricultural surveys. Hence, the ability to rapidly, remotely and cost-effectively quantify the damage to agricultural production area over large spatial extents can be an invaluable tool.

In September 2017, within two weeks of each other, hurricanes Irma and Maria grazed the northern coast of the Dominican Republic causing widespread damage (IFRC, 2018). Irma hit first, passing the coast of the Dominican Republic on 7th September 2017, causing storm surges, wind damage and flooding. Irma maintained a 60 hour period of category 5 intensity, the second longest period on record (Blake, 2018). On 21^st^ September 2017, Maria passed the northern and eastern coasts at Category 3, bringing strong winds and heavy rain (Blake 2018). The country’s northern provinces of Esapillat, Monte Cristi, Puerto Plata, Santiago, Samana and Valverde were the worst affected areas (IFRC, 2018). Of these provinces, three — Monte Cristi, Santiago and Valverde — comprise the Dominican Republic’s main banana growing area (Espinal, 2015).

Bananas are one of the Dominican Republic’s most important agricultural products, as most of what is produced is exported (Raynolds, 2008). It is the world’s largest producer of organic bananas, and the 23^rd^ largest producer of bananas (Lernoud et al., 2017). The country has an estimated 27,000 ha of banana production with 16,000 ha cultivating bananas for export to the key markets of Europe and the USA (Espinal, 2015). The banana sector employs an estimated 32,000 people in the Dominican Republic (ILO, 2015). Uniquely for a large export focused banana producing country, production has a large smallholder component with around 2000 small farms, each covering less than 7.5 ha (BAM, 2016). As such, banana production contributes substantially to the country’s local and national economy. Export banana production is concentrated in the North West Line regions of Valverde (31%) and Monte Cristi (38%) (Espinal, 2015). The provinces are dominated by the drainage basin of the Yaque del Norte river which runs through the Cibao valley. Here the rivers flood plains are vital for agricultural production, for both domestic and export markets (World Bank, 2018). The river provides water for irrigation for key agricultural crops including rice and banana. The drainage basin has suffered from severe deforestation over the past years and this has affected the hydrological regime, further exasperating the scale of floods at times of heavy rain, and reducing the available water in the river at times of drought (World Bank, 2018). Heavy rain during the two consecutive hurricane events in 2017 led to soil saturation, increased runoff in the drainage basin, and eventually the Yaque del Norte bursting its banks in several places (IICA, 2017). In addition, tributaries and drainage canals also overflowed.

Bananas are a semi-perennial crop, with the exportable Cavendish variety requiring approximately 9-10 months from planting to first harvest. Thereafter, plants enter a continuous production and harvest cycle (Heslop-Harrison & Schwarzacher, 2007). This implies a considerable lag (and knock-on economic consequences) between the loss of production following flood-or storm-related damage, requiring replacement of plants, and a return to previous production capacity. Consequently, hurricane impacts to banana growing regions of the Dominican Republic have food security and economic consequences, making evaluation of the damage important.

Earth observation or satellite remote sensing data is well suited for quantifying the impacts of extreme weather events (Sanyal & Lu, 2004; Plank, 2014; de Beurs, McThompson, Owsley, & Henebry, 2019). Data from Synthetic Aperture Radar (SAR) sensors, such as from the European Space Agency’s (ESA) Sentinel-1 platform, are particularly useful in detecting structural changes on the Earth’s surface, and hence have been widely used in natural disaster mapping and monitoring (Plank, 2014). Detecting open water in a landscape is made relatively easy when using SAR data, as the surface of water displays characteristically low reflection (backscatter) of the radar signal back to satellite-borne sensors (Schumann & Baldassarre, 2010; Twele, Cao, Plank, & Martinis, 2016). Hence, flooded areas are readily detectable by comparing SAR data for a location immediately before and after a storm or flooding event.

There is a long history of research using satellite data for land-use cover mapping (Townshend, Justice, Li, Gurney, & McManus, 1991; Defries & Townshend, 1994; Tuanmu & Jetz, 2014; Joshi et al., 2016), including the delineation of crop types in a landscape (Jansenn & Middlekoop, 1992; Inglada et al., 2015). Until recently, such mapping has largely relied on the analysis of multi-spectral satellite imagery (Jansenn & Middlekoop, 1992; Li, Wang, Zhang, & Lu, 2015). As SAR data (which has only recently become more widely available) conveys a measure of crop canopy structure or texture, its inclusion in such mapping methods adds an extra dimension of information that could increase mapping and classification accuracy (Inglada, Vincent, Arias, & Marais-Sicre, 2016). An additional advantage of incorporating SAR data into crop mapping is the ability to leverage temporal information of the backscatter signature. As SAR data from satellite platforms are not affected by cloud cover (as multi-spectral sensors are), an uninterrupted time series of SAR imagery over a landscape of interest can provide temporal parameters, such as periodicity and variance over time. Such metrics can be very powerful in the separation of crop classes, for example annuals from perennials, or amongst the annuals, summer and winter crops (Inglada et al., 2016; Veloso et al., 2017). These properties of SAR data could prove advantageous for the mapping of banana plantations, as (a) banana plants have a characteristic upright stature with large leaves, which we expect to result in high backscatter in SAR data; (b) they are perennial; and (c) commercial banana plantations (especially those catering to the export market) usually operate under a continuous production system (i.e. plants are infrequently replanted), and hence should show low variation in backscatter over time (e.g. in a year) compared to other crop types and vegetation classes.

Here we quantify the area of banana production in the Dominican Republic impacted by hurricanes Irma and Maria in September 2017, and focus on the Monte Cristi, Santiago and Valverde provinces, where the majority of the country’s production is concentrated. Specifically, we aim to (1) map flooded area, or more generally, area impacted by hurricane damage in the three provinces of interest using ESA’s Sentinel-1 data; (2) develop a classification algorithm to map commercial banana plantations using a combination of Sentinel-1, Sentinel-2, and topographic data using post-hurricane ground truth data of banana plantations; (3) apply the banana plantation classification method to pre-hurricane satellite data in order to quantify area of plantations affected by Irma and Maria in 2017; and (4) identify areas of current banana plantations in the study region at risk from similar extreme weather events. In the process, we also aim to design a classification method to map commercial plantations that could be more widely applied to banana production systems globally.

## Methods

Our study area includes the provinces of Monte Cristi, Santiago and Valverde in the Dominican Republic. For reference, hurricanes Irma and Maria struck the region on the 7th and 21st of September, 2017, respectively. Hence, we label satellite data before this date range as pre-hurricane, and after this date range as post-hurricane.

### Mapping hurricane damage

We primarily rely on ESA’s Sentinel-1 data which we processed using Google Earth Engine to map hurricane damage. The total extent of damage was mapped as three separate components which were then combined. First, detectable open water flooding immediately after each hurricane was classified. We term this component ‘flood-open’ (FO). Second, an arbitrary fixed distance around each FO patch (100 meters) was also hypothesised to experience flood damage. This was done because tall vegetation elements, such as banana plants, which may not have been immediately damaged during the hurricane could, in part, obscure the signal of open water in the SAR data. However, it is likely that the ground in these obscured pixels would have been inundated (Figure 1), or saturated enough to stress any standing crop. This component was termed ‘flood-buffer’ (FB). Third, we mapped ‘flood-legacy’ (FL) as pixels which display large deviations (described in detail below) in the three months after the hurricanes, relative to time-averaged pixel values for a year prior to the hurricanes. This latter component accounts for more protracted or delayed damage following the flooding events. We estimate the total hurricane affected area for the study region as the spatial union of the three components (i.e. FO ∪ FB ∪ FL), and provide detailed methods for FB ∪ FB ∪ FL), and provide detailed methods for FL), and provide detailed methods for the mapping of each component below.

**Figure 1.**
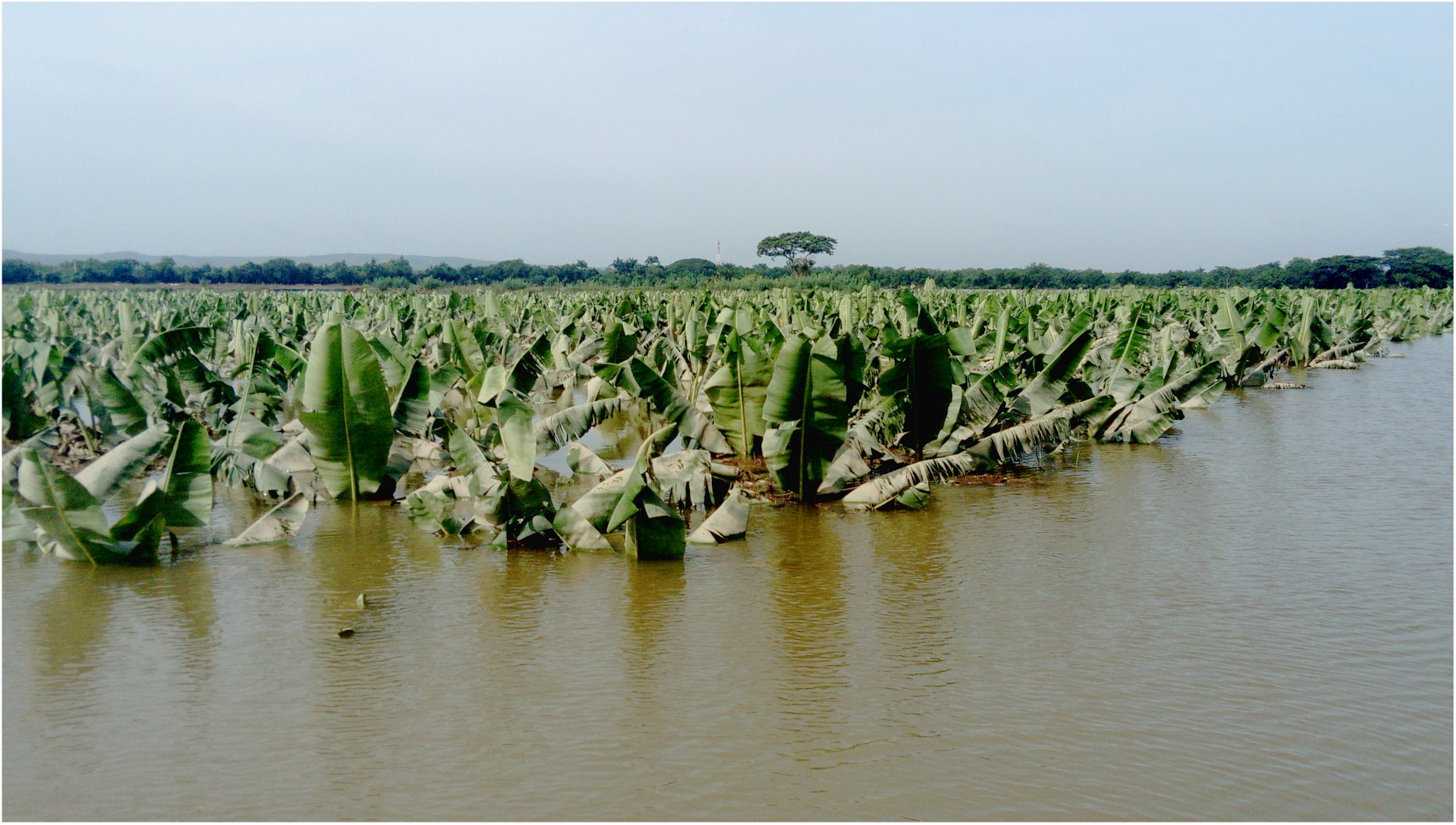
Extensive flooding in a banana plantation in the Dominican Republic in September 2017. The image shows that some open water flooding could be obscured by the canopy of standing banana plants *[image credit: annon]*.

For the FO component of flooding, we used SAR imagery for the three provinces immediately before (B) hurricane Irma, immediately after hurricane Irma (I) and immediately after hurricane Maria (M). We applied median smoothing using a circular kernel of 100 meter radius to each image, to reduce speckling noise that SAR data from a single time snapshot can suffer from. Thereafter, we calculated the pixel-wise differences between I and B (flood damage due to Irma), and M and B (flood damage due to Maria). We assumed that flood or storm related structural damage to landscape elements reduces backscatter in the SAR data. Additionally, pixels which contain open water after a flood event would also show considerably reduced backscatter compared to values when the pixel was not flooded, i.e. prior to the flooding event (Schumann & Baldassarre, 2010). Hence, we identified pixels in the difference data layers which showed values < −2 dB to have experienced flood or storm damage. A separate binary map of these affected regions was generated for hurricanes Irma and Maria. Pixels which fell within 100 meters from these FO pixels were categorised as FB pixels, i.e likely to have experienced soils saturated by moisture, if not inundated by flood water.

To map flood-legacy (FL), we extracted all Sentinel-1 images for a one year period prior to the hurricanes (1st September, 2016 to 6th September, 2017) — before image set (BS), and three months during and after the hurricanes (6th September, 2017 to 30th November, 2017) — after image set (AS). We only utilised the VV polarisation of the Sentinel-1 data for our analyses, as many images, especially those in 2016, do not contain the VH polarisation band. The 26 images from the BS subset were used to calculate the annual pre-hurricane average VV backscatter for the region (BS_mean_). Similarly, we calculated the standard deviation for each pixel using this annual stack of images (BS_SD_). The post-hurricane average VV backscatter was calculated for the AS subset using its seven images (AS_mean_). Next, the amount of the deviation in mean VV values between the pre-and post-hurricane data subsets was calculated in units of standard deviations D = [(AS_mean_ - BS_mean_) / BS_SD_]. All pixels below an arbitrary threshold of D ≤ −0.5 SD were retained as candidate pixels displaying flood-legacy effects (here again we assume loss of structural complexity in the wake of storm damage resulting in lower backscatter intensity). To exclude single pixel artefacts, a 7×7 (i.e. 70 meter x 70 meter) moving window was used to count the number of candidate pixels around each focal pixel. Each focal pixel within a window was reassigned as having experienced flood-legacy effects if the number of candidate pixels within the window were ≥ 20 (i.e. approximately 40%). This is an arbitrary threshold which we believe provides a conservative estimate for the affected area. It is important to note that this FL component is not calculated separately for hurricanes Irma and Maria as was the case with the FO and FB components.

### Banana plantation area mapping

Ground truth data of 100 banana farm polygon boundaries for classifying and validating banana plantation maps was sourced from ground surveys of farms conducted by *Bananos Ecológicos de la Línea Noroeste* (BANELINO), a producer co-operative, in 2019. Additional land-cover classes were also included in our analysis by manually digitising training classes from images available on Google Earth. These additional classes were urban/semi-urban/rural built area, other crops (crops that were visually distinguishable from bananas), mangroves, dense natural/semi-natural tree cover and sparse natural/semi-natural tree cover.

We extracted Sentinel-1 and Sentinel-2 data of the date range corresponding to the ground truth data in Google Earth Engine for processing. As the commercial cultivation of bananas is conducted on relatively flat terrain, slope derived from 90 meter resolution Shuttle Radar Telemetry Mission (SRTM) digital elevation model (DEM) was also included in the classification procedures.

Sentinel-1 data from one year that spanned the time period that best corresponded with the ground-truth data was extracted. The 26 images from this period were aggregated to two derived layers - annual median VV and annual standard deviation of the VV bands. The surface reflectance product of Sentinel-2 (all images for the same time period as the Sentinel-1 subset) was used to calculate a median red, green, blue and NDVI layer. These derived bands were calculated following masking of pixels within each individual Sentinel-2 image for cloud contamination based on the quality assessment band for each individual image.

The two derived layers from Sentinel-1 imagery, four derived layers from Sentinel-2 imagery and the slope layer were stacked to form a single multi-band image, and used to build the classifier. Five hundred random sampling points within polygons of each land-cover class were generated. These were used to build a random forest classifier using 50 decision trees. To increase accuracy and reduce classification artefacts, pixels classified as banana were only retained if they formed a patch of ≥ 50 pixels (i.e. ≥ 0.5 ha). Hence, it is important to note that the method described here is not suitable to detect small plantations or mixed cropping.

Accuracy of the classifier was estimated by generating a separate set of 500 random sampling points within each land-cover class and running the classifier over this test dataset. A confusion matrix (Stehman, 1997) was then constructed to estimate training and validation accuracy.

To map pre-hurricane banana production area, we processed Sentinel-1 and Sentinel-2 imagery for one year before the hurricanes (i.e. 1^st^ September, 2016 to 30^th^ August, 2017) in a similar way to that described above (the same DEM data was used to derive a slope layer). The random forest classifier was then used to generate a raster of banana production area distribution.

Raster layers of the three flooding components and pre-hurricane banana production distribution were exported to QGIS 3.10 (QGIS.org, 2019) and GRASS 7.4 (GRASS Development Team, 2018) for further processing. The flood components were combined into a single layer delineating total hurricane affected area. This layer was intersected with the pre-hurricane classified banana layer to measure the area of production affected by hurricane damage. We intersected the total hurricane affected area with the post-hurricane banana map to estimate the area of current banana production which may be at risk from damage should a similar extreme event recur. Lastly, we mapped the spatial turnover of plantation area between 2017 (pre-hurricane) and 2019 (current), which captures the spatial change in plantation distribution, i.e. where plantation area has been lost, and where there has been a gain. By overlaying the hurricane affected area data with areas of production area gain in 2019, we then quantified how much of the newly planted area could be at risk from damage from future storm damage.

## Results

A total of 27,618 ha of hurricane-related flood damage was estimated for the three provinces of Monte Cristi, Valverde and Santiago, equating to 5.01% of the total area of the provinces (Figure 2). Individually, flood-open (FO), flood-buffer (FB) and flood-legacy (FL) accounted for 8,302.3 ha, 18,178.21 ha and 15,622.9 ha, respectively. Total affected area is less than the sum of the three flood classes due to considerable spatial overlap between them. Observed flood damage was proportionally greater in Monte Cristi (18,102 ha; 9.3%) and Valverde (6,051 ha; 7.6%) compared to Santiago (3,462 ha; 1.3%). Additionally, flood damage in Monte Cristi and Valverde appeared to be more spatially aggregated around the Yaque del Norte river with large areas of open flooding. In contrast, Santiago saw more scattered occurrences of damage largely classified as legacy effects.

**Figure 2.**
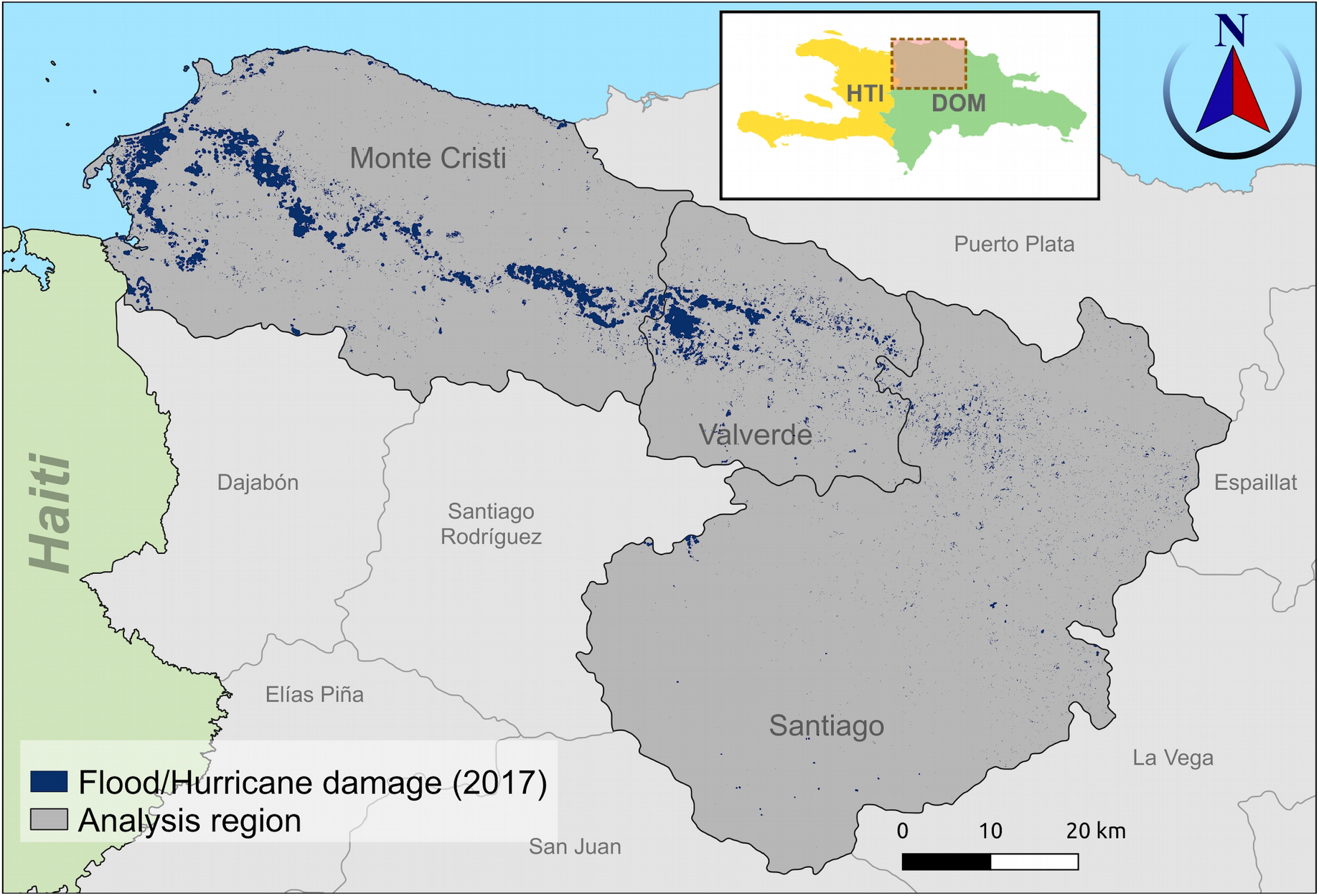
Extent of estimated hurricane damage in the provinces of Monte Cristi, Valverde and Santiago. Inset illustrates the region of analysis in the Dominican Republic.

The random forest classifier performed particularly well for mapping banana plantations in the three provinces. Overall accuracy estimated using a confusion matrix was 99.7% for all classes and 99.8% for banana plantations. In total, for 2019, we detected 23,898 ha of plantation area in the three provinces (Figure 3). Projecting the classifier on satellite images before hurricanes Irma and Maria in September 2017, we detected a total plantation area of 21,561 ha within the region of analysis (Figure 4).

**Figure 3.**
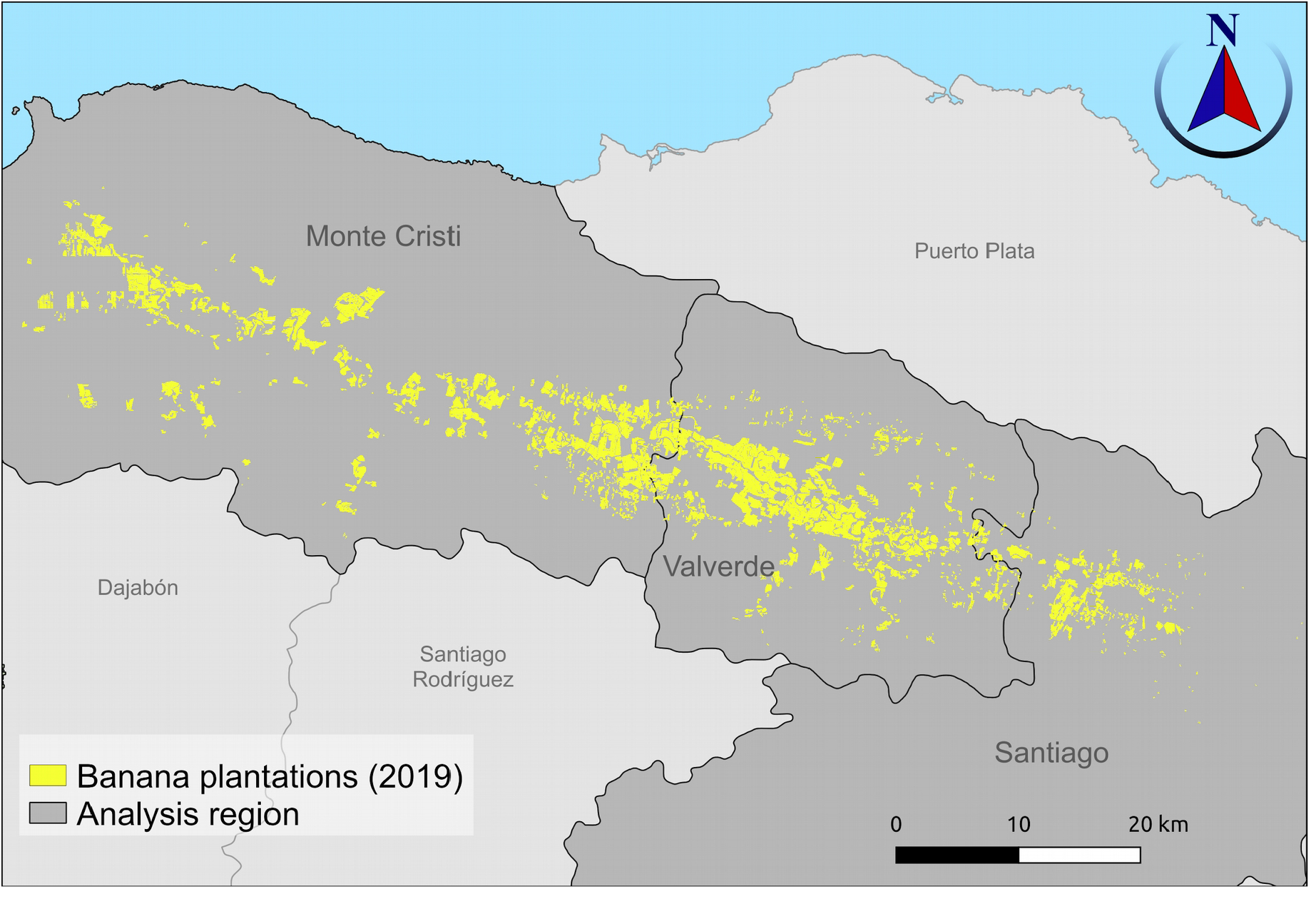
Distribution of banana plantation area in the study region in 2019 (post-hurricane)

**Figure 4.**
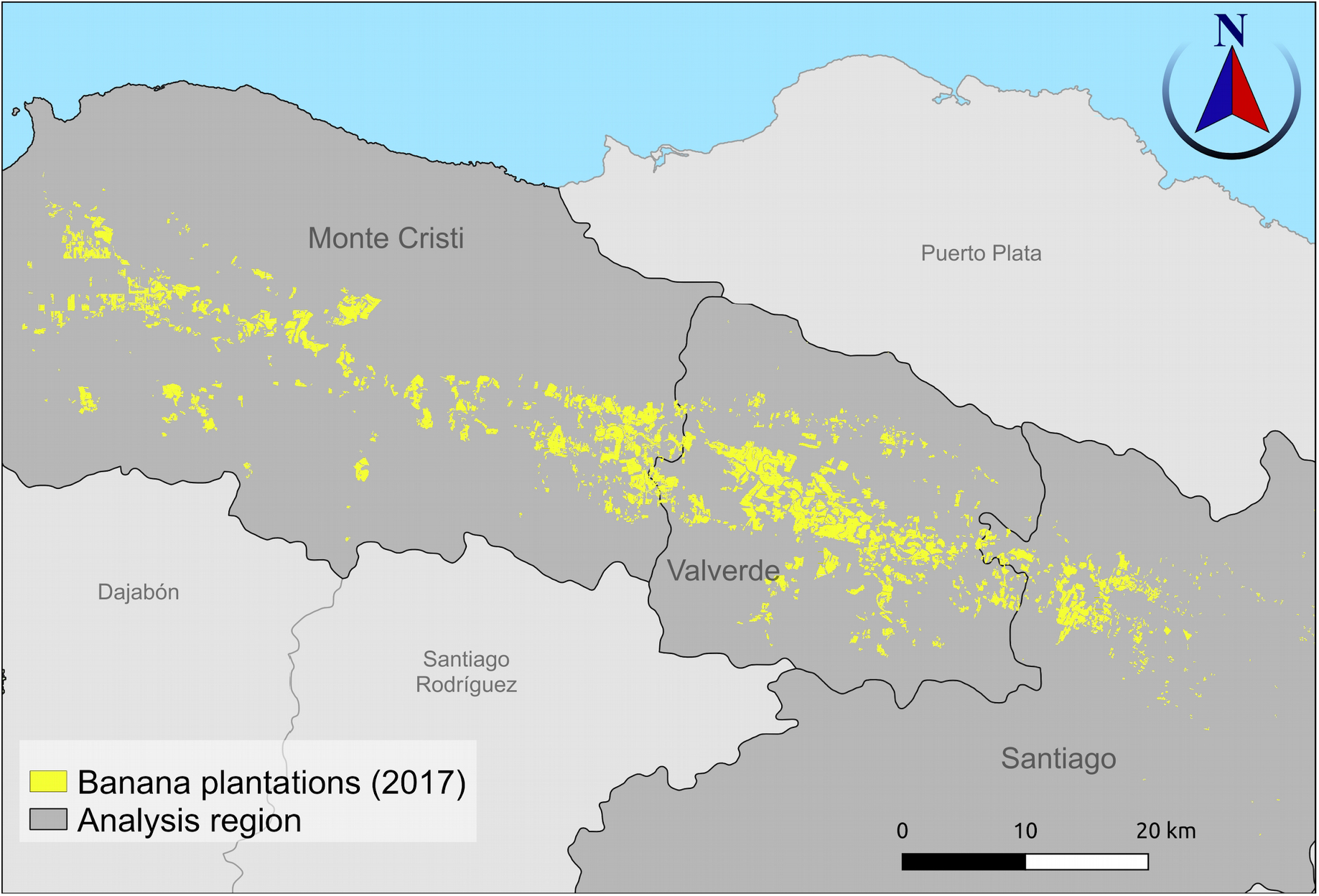
Distribution of banana plantation area in the study region before September 2017 (pre-hurricane plantation distribution)

On overlaying the hurricane damage (Figure 2) and banana plantation maps for 2017 (Figure 4), we estimate 2,446.75 ha of plantations were likely to have experienced damage. This accounts for 11.35% of area under cultivation before the hurricanes struck the Dominican Republic (Figure 5). By overlaying the hurricane damage map and banana plantation map for 2019 (Figure 6), we estimate 3,402.55 ha or 14.24% of plantation area may be at risk from damage should a similar extreme weather event reoccur.

**Figure 5.**
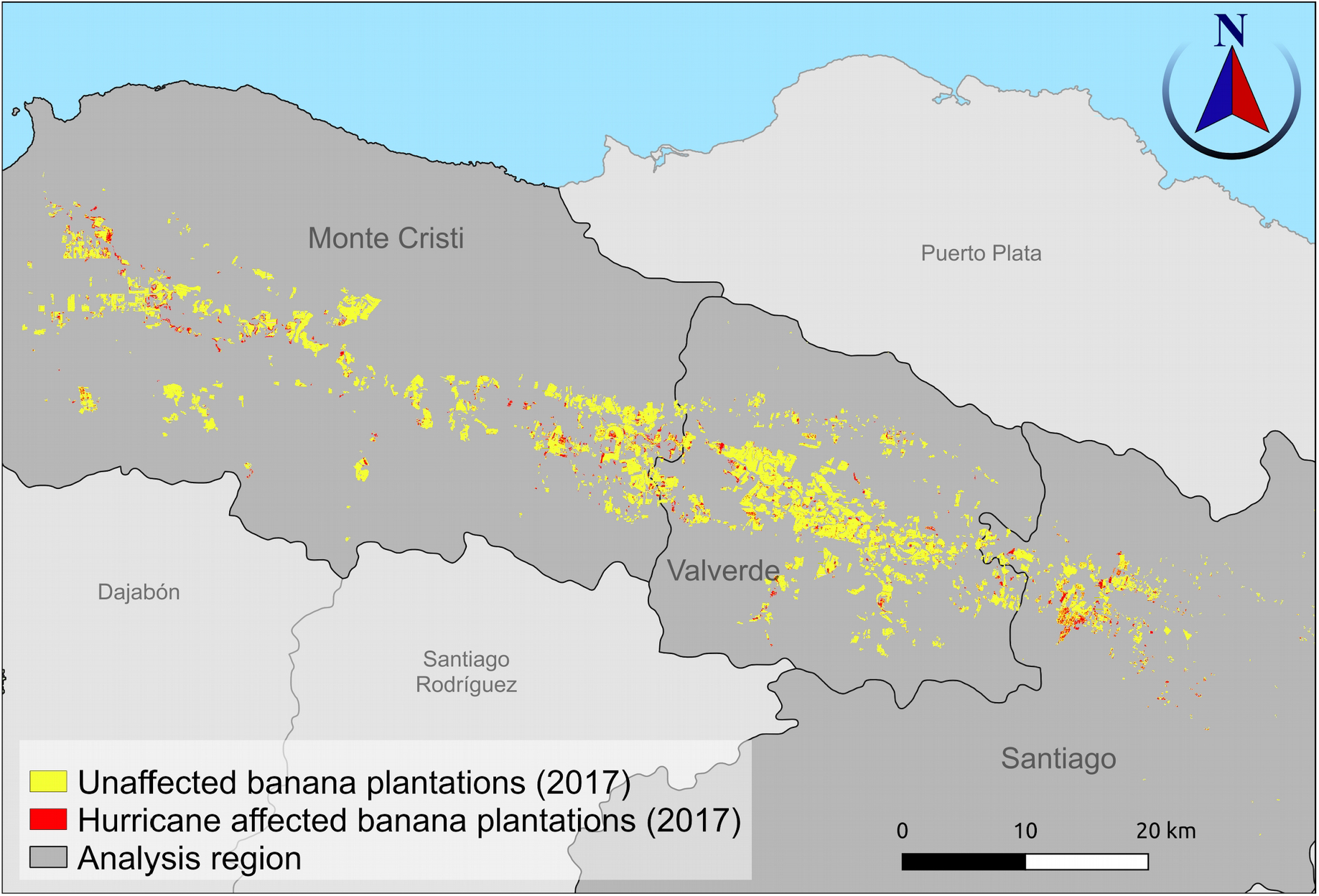
Map identifying locations in the study region where banana plantations in 2017 were affected by hurricanes Maria and Irma (September 2017)

**Figure 6.**
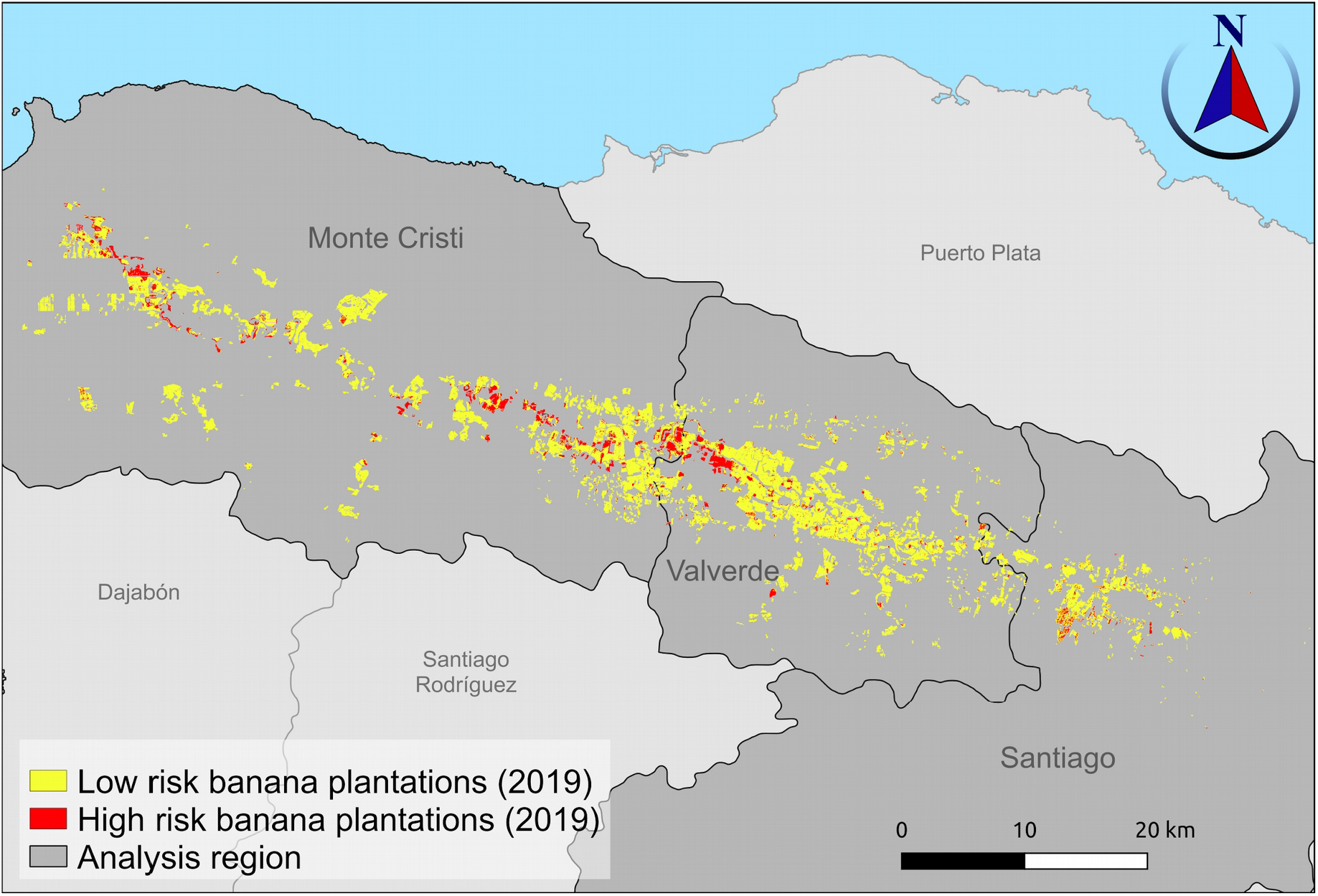
Banana plantation areas in 2019 that overlap with locations that saw storm-related damage due to hurricanes Irma and Maria in 2017. The highlighted regions (red) are considered at high risk should similar extreme weather events reoccur.

Our analyses also showed a loss of 5,048.37 ha of pre-hurricane banana plantation area (Figure 7). This amounts to 23.4% of the production area in 2017. Of the plantation area lost between 2017 and 2019, 1,031.41 ha was identified to have experienced storm damage. However, by 2019 there was also a gain of 7,384.84 ha of new plantation area. Importantly, 1987.21 ha or 26.9% of this new plantation area overlapped with locations which saw damage in the aftermath of hurricanes Irma and Maria.

**Figure 7.**
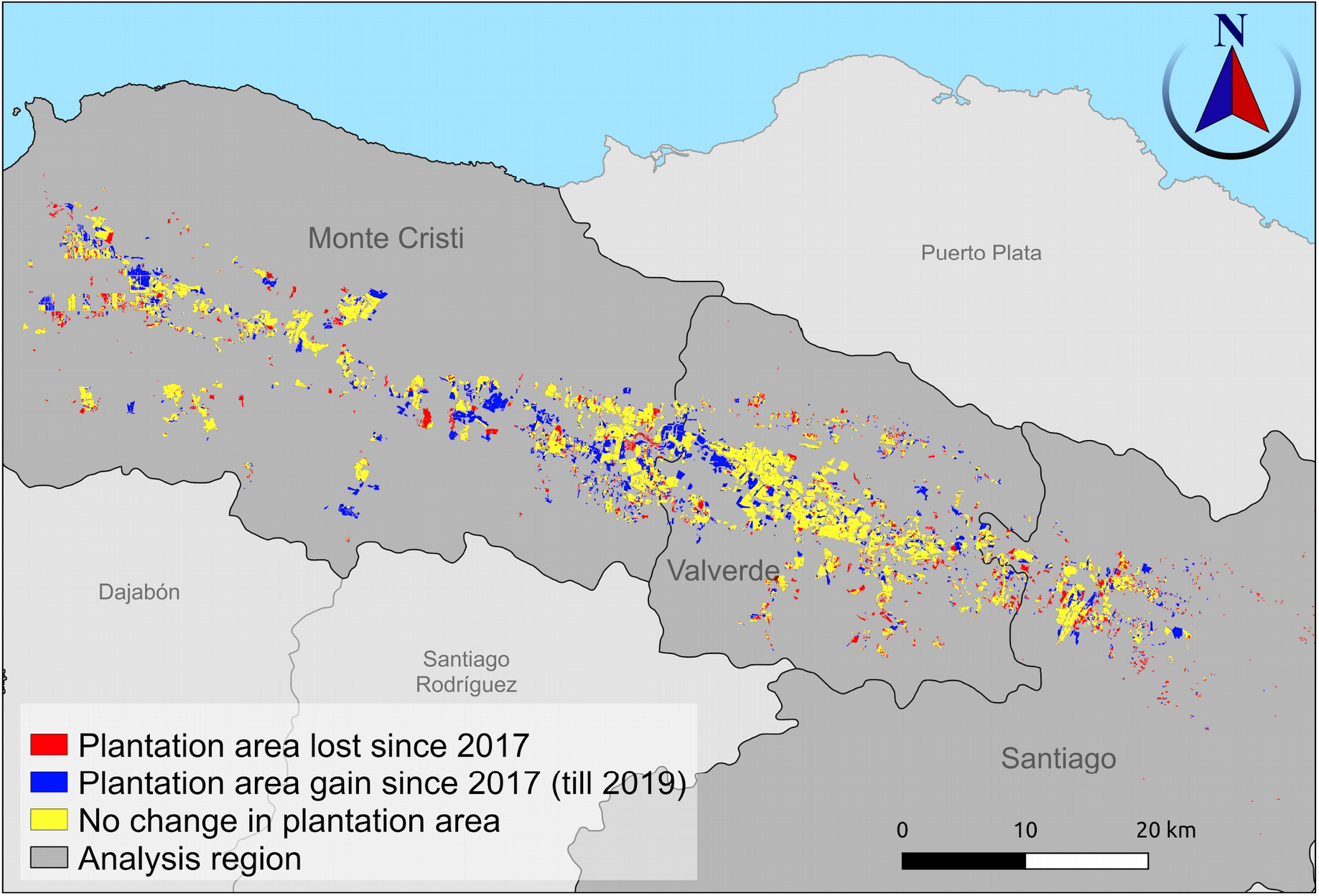
Map illustrating plantation area turnover in the study region between 2017 and 2019.

## Discussion

Damage from hurricanes Irma and Maria was experienced in all the three provinces of Monte Cristi, Valverde and Santiago, though the area affected in the latter was considerably less, and spatially more dispersed. Overlaying the hurricane damage map with our classified pre-hurricane banana plantation map we estimate that 2,446.75 ha, or 11.35% of banana plantation area across the three provinces was affected by open water and/or protracted storm damage over a period of three months since the hurricanes. The damage to plantations was spatially more evenly spread across the three provinces.

These results reveal large scale damage to a key export sector of the Dominican Republic. Uncertainty around adequate production following the hurricanes led to importers of large European retailers switching procurement to other countries. To alleviate the economic consequences to farmers through the loss of market access — even for those whose production area may have been largely unaffected — the national government had to intervene by procuring production initially destined for the export market, thereby severely affecting the country’s economy (FAO, 2018; Polanco, 2018). Rapid assessment of the scale of damage, as carried out in this study, could allow importers to maintain sourcing in the Dominican Republic by providing more accurate and up-to-date information to base sourcing decisions on.

Our analyses estimated that 23.4% (5,048.37 ha) of area under banana cultivation before hurricanes Irma and Maria had been converted to non-banana producing area by 2019. The converted area is much larger than what we estimated as having been affected by the two hurricanes. Additionally, only 42.2% of the banana plantation area of 2017 that showed evidence of storm damage was converted to other land-use classes. Hence, we conclude that losses in banana cultivation area between 2017 and 2019 may have only partly been driven by direct hurricane damage, and that other local economic factors — which may include indirect consequences of hurricane impacts — may have been equally as important and require further investigation.

Despite the localised losses of production area between 2017 and 2019, the area under banana cultivation across the study area grew by 10.8% with the addition of 7,383.84 ha of new plantations. Over a quarter (26.9%; 1987.21 ha) of this new production area overlaps with areas that experienced hurricane related damage in 2017. While the incidence of two large magnitude hurricanes in rapid succession can be considered as rare, there have been consistent predictions that the frequency and intensity of such extreme events are likely to increase due to climate change (IPCC, 2014). Hence, we identify these areas (figure 6) as at risk. Further, this also results in 14.24% of the area under banana cultivation at risk from similar storm events in the future. For context, this is an increase from 11.35% of production area which saw damage in 2017. A comprehensive assessment, as presented here, could be used to inform risk management measures and risk transfer solutions, such as investment in micro-insurance. Additionally, risk maps could enable efficient aggregation of risk across co-operatives or administrative units spanning the production landscape.

While our analyses suggest that production area in the main banana growing areas of the Dominican Republic has shown a complete recovery since the hurricanes Irma and Maria, finer scale (i.e. farm level) patterns of recovery are yet to be investigated. Banana production in the Dominican Republic comprises a large number of smallholder farmers, and there is wide consensus in the literature that smallholders are at particular risk from climate change and associated extreme events (Morton, 2007; Harvey et al., 2014). Future research should focus on linking satellite derived plantation-scale data at high temporal resolution with information on farm-scale economic status and decisions, as well as production volumes. Such models could be important for formulating wider government-led recovery strategies, and informing importer procurement decisions.

We obtained high levels of accuracy for our classification method to map banana plantations using a combination of Sentinel-1, Sentinel-2 and elevation data. This gave us a high degree of confidence in mapping pre-hurricane plantation distribution in the study area, and consequently, in estimating the extent of hurricane related damage to plantations. Bearing in mind that the random forest classifier was trained only using 500 sampling points for the banana class (i.e. approximately 5 ha of plantation area), the high accuracy suggests that even with relatively low effort in ground-truthing, this method represents a very promising approach to mapping commercial banana plantations more widely. Bananas are one of the most extensively cultivated crops in the world (FAOSTAT, 2020). Like any other commercially cultivated crop, banana production also faces challenges from multiple stressors, apart from extreme weather events, such as longer-term climate shifts, pests and diseases (Ramirez, Jarvis, Van den Bergh, Staver, & Turner, 2011; Ordonez et al., 2015; Bebber, 2019; García-Bastidas et al., 2019; Varma & Bebber, 2019). Assessing risk from these stresses, measuring impacts, as well as monitoring rates of recovery, and effectiveness of mitigation measures require detailed information on plantation distribution. Such spatially explicit data can also be invaluable in tracking the performance of plantations, by providing focal areas over which to analyse finer variations in measurements from satellite-borne sensors. However, such up-to-date and high resolution information on the distribution of banana plantations is lacking, and methods described here could help address this gap.

In conclusion, this study used remote sensing data and analyses to provide a detailed assessment of the impacts of hurricanes Irma and Maria on production area of bananas in the key growing areas of the Dominican Republic. With the open availability of regularly captured data from satellite platforms, such as ESA’s Sentinel-1 and Sentinel-2, assessments such as these soon after the occurrence of an extreme weather event can be rapidly and cost effectively carried out. Our analyses also mapped where current production area may be at risk from similar high intensity storm events in the future. We observed that there has been a net increase in banana plantation area in the region between 2017 and 2019, a substantial proportion of which has occurred in locations which experienced hurricane related damage in 2017. Consequently, there has been an overall increase in production area at risk from future storm events which should be considered in mitigation and recovery strategies. Lastly, we have demonstrated a highly accurate method to map banana plantation area that can form the basis of tracking the trajectory of banana production recovery from extreme weather events. Research focused on quantifying these patterns of recovery could positively contribute to government risk management and mitigation planning, importer procurement decision making, and ultimately, securing farmer livelihoods.

## Funding information

This study was funded by Global Food Security grant no. BB/N020847/1; EC Horizon 2020 project ID 727624; Science and Technology Facilities Council (STFC) Food Network; Mercator Research Program of the ETH Zurich World Food System Centre and the ETH Zurich Foundation.

## Abbreviations used

AS: ‘After set’ of images; A three month collection of Sentinel-1 images (seven images) during and after hurricanes Irma and Maria affected the study area (6th September 2017 to 30th November 2017).
AS_mean_: A single band image of mean pixel values from AS.
B: Sentinel-1 image over the study region immediately before hurricane Irma.
BS: ‘Before set’ of images; A one year collection of Sentinel-1 images (26 images) before hurricane Irma affected the study area (1st September 2016 to 6th September 2017).
BS_mean_: A single band image of mean pixel values from BS.
BSSD: A single band image of pixel wise standard deviations from BS.
D: A single band image of the difference between AS_mean_ and BS_mean_ expressed in terms of BSSD
DEM: Digital Elevation Model
ESA: European Space Agency
FB: Flood-Buffer; regions upto 100m from pixels detected as open-water flooding (FO)
FL: Flood-Legacy; Pixels assessed to have experienced more protracted hurricane/flood damage. They are characterised by large deviations in pixel values in the three months following hurricanes Irma and Maria, relative to values observed for the same pixel over a one year period before the hurricanes.
FO: Flood-Open; regions which show characteristics of open-water flooding in Synthetic Aperture Radar satellite data
I: Sentinel-1 image over the study region immediately after hurricane Irma.
M: Sentinel-1 image over the study region immediately after hurricane Maria.
NDVI: Normalised Difference Vegetation Index
SAR: Synthetic Aperture Radar
VH polarisation: Vertical transmit - horizontal receive band of Sentinel-1 imagery
VV polarisation: Vertical transmit - vertical receive band of Sentinel-1 imagery

## Notes

### Competing Interest Statement

The authors have declared no competing interest.

